# A tiger in the Upper Midwest: Surveillance and genetic data support the introduction and establishment of *Aedes albopictus* in Iowa, USA

**DOI:** 10.1101/2021.10.20.465182

**Authors:** David R. Hall, Ryan E. Tokarz, Eleanor N. Field, Ryan C. Smith

## Abstract

*Aedes albopictus* is a competent vector of several arboviruses that has spread throughout the United States over the last three decades after it was initially detected in Texas in 1985. With the emergence of Zika virus in the Americas in 2015-2016 and an increased need to better understand the current distributions of *Ae. albopictus* in the US, we initiated surveillance efforts to determine the abundance of invasive *Aedes* species in Iowa. Here, we describe the resulting surveillance efforts from 2016-2020 in which we detect stable and persistent populations of *Aedes albopictus* in three Iowa counties. Based on temporal patterns in abundance and genetic analysis of mitochondrial DNA haplotypes between years, our data support that populations of *Ae. albopictus* are overwintering and have likely become established in the state. In addition, the localization of *Ae. albopictus* predominantly in areas of urbanization and noticeable absence in rural areas suggests that these ecological factors may represent potential barriers to their further spread and contribute to overwintering success. Together, these data document the establishment of *Ae. albopictus* in Iowa and their expansion into the Upper Midwest, where freezing winter temperatures were previously believed to limit their spread. With increasing globalization, urbanization, and rising temperatures associated with global warming, the range of invasive arthropod vectors, such as *Ae. albopictus*, is expected to only further expand, creating increased risks for vector-borne disease.

## Introduction

*Aedes albopictus* is an invasive mosquito species that in recent decades has spread across multiple continents predominantly through global trade [1–4]. With the first report of its establishment in the United States (US) in Texas in 1985 [5], its range has continually expanded to more than 26 states, gradually spreading northward and westward across the US [6–8] with likely further expansion fueled by climate change and increasing urbanization [3]. With anthropophilic feeding behavior [4,9] and as a competent vector of at least 26 mosquito-borne arboviruses [10], the introduction of *Ae. albopictus* into new locations in the US raises a significant public health concern.

With *Ae. albopictus* as a competent vector of Zika virus (ZIKV) [11,12], the emergence of ZIKV in the Americas in 2015 and 2016 created a critical need to better understand the distributions of *Ae. albopictus* in the US in order to determine the potential risks for ZIKV transmission. Previous studies have described the detection of *Ae. albopictus* populations across the Midwest [7,8,13–17]. This includes Iowa, which has seen sporadic detections of *Ae. albopictus* that likely represent rare and unsuccessful introduction events [17]. However, with established populations of *Ae. albopictus* in the neighboring states of Missouri [7,8,13] and Illinois [7,8,16], there is a high likelihood of further *Ae. albopictus* introduction events and potential invasion with modeling suggesting that Iowa is within this species’ predicted range [18]. Initial targeted surveillance in 2016 along the southern Iowa border failed to detect *Ae. albopictus* [19], yet with sampling only over a single year, there may not have been adequate trapping efforts to identify low-density populations.

In this study, we describe our continued monitoring of mosquito populations in Iowa through targeted surveillance efforts focusing on invasive *Aedes* species. Expanding on our initial efforts [19], we used a trapping network consisting of BG sentinel and Gravid *Aedes* traps from 2017 to 2020 to monitor mosquito populations in a total of 25 counties over a five year period (2016 to 2020). Through these efforts, we document the detection and likely establishment of *Ae. albopictus* in three Iowa counties. In addition, we provide evidence of their intra-county movement and the ecological variables that define their presence and absence, elucidating the potential ecological barriers that have thus far prevented their further spread to adjacent counties. Genetic analysis confirms the subsistence of genetic haplotypes between years, supporting the establishment of *Ae. albopictus* in each of the respective counties in which it has been detected, providing insight into the origins of their introduction.

## Methods

### Mosquito Surveillance

Targeted mosquito trapping efforts were performed by Iowa State University personnel or in collaboration with local public health departments between 2016 and 2020 from mid-May through October when mosquitoes are most active in Iowa. While initial efforts in 2016 relied on the use of BG Sentinel and CDC light traps [19], trapping from 2017-2020 utilized BG Sentinel 2 (BG) traps and Gravid *Aedes* Traps (GATs) (Biogents, Regensburg, Germany). These traps have proven to be highly effective in capturing *Aedes albopictus* and other container-breeding *Stegomyia* specimens [20–24], with the relatively low cost and little maintenance of the GATs enabling broader coverage with each respective county. BG traps were used without a carbon dioxide source, relying on human scent lures (BG-Lure, Biogents), while the GATs were equipped with sticky cards to enable mosquito collections.

Following our efforts in 2016 which targeted 9 of the 10 counties along Iowa’s southern border [19], we expanded our trapping efforts in 2017 to incorporate all eastern border counties along the Mississippi River and counties with more densely populated cities. These counties were chosen based on their proximity to Missouri and Illinois which have established populations of *Ae. albopictus* [7,8,13,16] and that have the highest potential for introduction via shipping/transport into more densely populated areas. In subsequent years (2018-2020), data from the previous trapping efforts allowed for more focused surveillance, reducing the overall number of participating counties. Trapping efforts were continued in each county where *Ae. albopictus* was detected, as well as in counties that represented potential sites of introduction or spread from adjacent positive counties.

### Mosquito Sample Processing and Identification

Mosquito samples were collected three times a week from BG Sentinel traps, while GATs were collected on a weekly basis. Samples were either transported directly or shipped to Iowa State where mosquitoes were identified to species using morphological characteristics and taxonomic keys [25]. Corresponding data were recorded according to date, trap location, and trap type. *Ae. albopictus* samples were separated into site/date-specific vials and stored in an ultra-low temperature freezer at -80°C for later genetic analysis.

### Geographic and Land Cover Analysis

The latitude and longitude coordinates of all trap locations were recorded and utilized to plot trapping locations using QGIS version 3.14.1. A Landsat-based, 30-meter resolution land cover layer, clipped to reflect our study area, was obtained from the Multi-Resolution Land Characteristics Consortium (MRLC) National Land Cover Database (NLCD) [26] which served as the base map and the source of all land cover output. Land cover analysis was performed for the two counties in southeast Iowa (Lee and Des Moines counties) which displayed widespread presence of *Ae. albopictus*. Trapping site locations within these counties were examined in QGIS using a 500 meters radius around each surveillance site, with the zonal histogram tool to list pixel counts of each unique land cover value within that radius (buffer layer). This distance was chosen based on the limited flight range of *Ae. albopictus*, which typically does not travel far from its site of origin with a maximum flight range of ∼500m [27–29]. Based on pixel numbers corresponding to each land cover feature, the percent land cover was calculated for each site and used to compare between locations where *Ae. albopictus* was present or absent.

### Genetic Analysis

*Ae. albopictus* samples collected from site locations between 2017 and 2018 in each of the three positive Iowa counties were examined by genetic analysis. A total of 165 samples were processed using the Marriott DNA extraction procedure [30–32] to isolate genomic DNA which was used as a template for PCR genotyping. Similar to other studies that have examined *Ae. albopictus* genetic haplotypes [16,33], a fragment of the mitochondrial cytochrome c oxidase subunit 1 was targeted with the following primers: 2027F (5’-CCC GTA TTA GCC GGA GCT AT-3’) and 2886R (5’-ATG GGG AAA GAA GGA GTT CG-3’). PCR was performed using DreamTaq Green DNA Polymerase (Thermo Fisher Scientific) under the following conditions: initial denaturation 94°C, 3 min; denaturation 94°C, 30 sec; annealing 55°C, 30 sec; extension 72°C, 1 min for 35 cycles; and a final extension 72°C, 6 min. PCR products were examined by electrophoresis on a 1% agarose gel, excised, and recovered using a Zymoclean Gel DNA Recovery Kit (Zymo Research). Resulting DNA was cloned into a pJET 1.2/blunt cloning vector using the CloneJET PCR Cloning Kit (Thermo Fisher Scientific), and subsequently transformed into DH5-alpha competent *E. coli* (New England Biolabs). Bacteria were plated on LB agar plates with a 100 μg/ml ampicillin concentration and incubated overnight at 37°C to select for successfully transformed colonies. Individual colonies were randomly chosen from the selection plates, suspended in 3 ml of LB broth and cultured overnight in a 37°C shaker at 215 RPM. Plasmid DNA was isolated using the GeneJet Plasmid Miniprep Kit (Thermo Fisher Scientific), with the presence of an insert validated by *Bgl*II digests and gel electrophoresis. Sanger sequencing of the resulting samples was conducted by the Iowa State University DNA Facility.

DNA sequencing data was aligned and edited manually using BioEdit version 7.0.5.3. To minimize the possibility of polymerase error, at least 3 sequences from each individual sample were combined to create a consensus sequence for each sample. Any unique sequences were confirmed by the additional amplification using Phusion High-Fidelity DNA Polymerase (Thermo Fisher Scientific) followed by cloning and sequencing using the above methods. DNA from individual mosquito samples were grouped into haplotypes where each haplotype represents a unique sequence, and the number of polymorphic sites, haplotype diversity (*Hd*), and nucleotide diversity (π) were calculated using DnaSP (version 6.12.03) [16,33,34]. A haplotype network was created in PopART [35] using the median–joining network method [36] to visualize genetic relationships between haplotypes and to display differences in population structure between sites [16].

## Results

### Mosquito surveillance and detection of *Ae. albopictus* in Iowa

To determine if the invasive mosquito species, *Ae. aegypti* and *Ae. albopictus*, could be found in the state of Iowa, we performed targeted mosquito surveillance in a total of 25 counties from 2016 to 2020 (Figure 1A, Table S1, Table S2). After initial surveillance efforts along the southern border of the state in 2016 [19], we extended our trapping efforts from 2017-2020 to more densely populated counties and to those bordering the Mississippi River (Figure 1A, Table S1, Table S2). Although *Ae. aegypti* and *Ae. albopictus* were not detected in 2016 [19], a total of 432 *Ae. albopictus* were collected in 2017 from three Iowa counties (Polk, Lee, and Des Moines) (Figure 1A and 1B). In subsequent years (2018 to 2020), *Ae. albopictus* were similarly detected in each of the same three counties in increased numbers, reaching a high of 1,315 *Ae. albopictus* detected in 2020 (Figure 1B). From 2017 to 2020, a total of 3,700 *Ae. albopictus* were collected, with Lee County displaying the highest total of *Ae. albopictus* amongst the three counties (Figure 1C) and consistently producing the highest number of *Ae. albopictus* between years (Figure 1D). Together, these data suggest that in recent years *Ae. albopictus* have been introduced into the state, and have potentially become established in three Iowa counties.

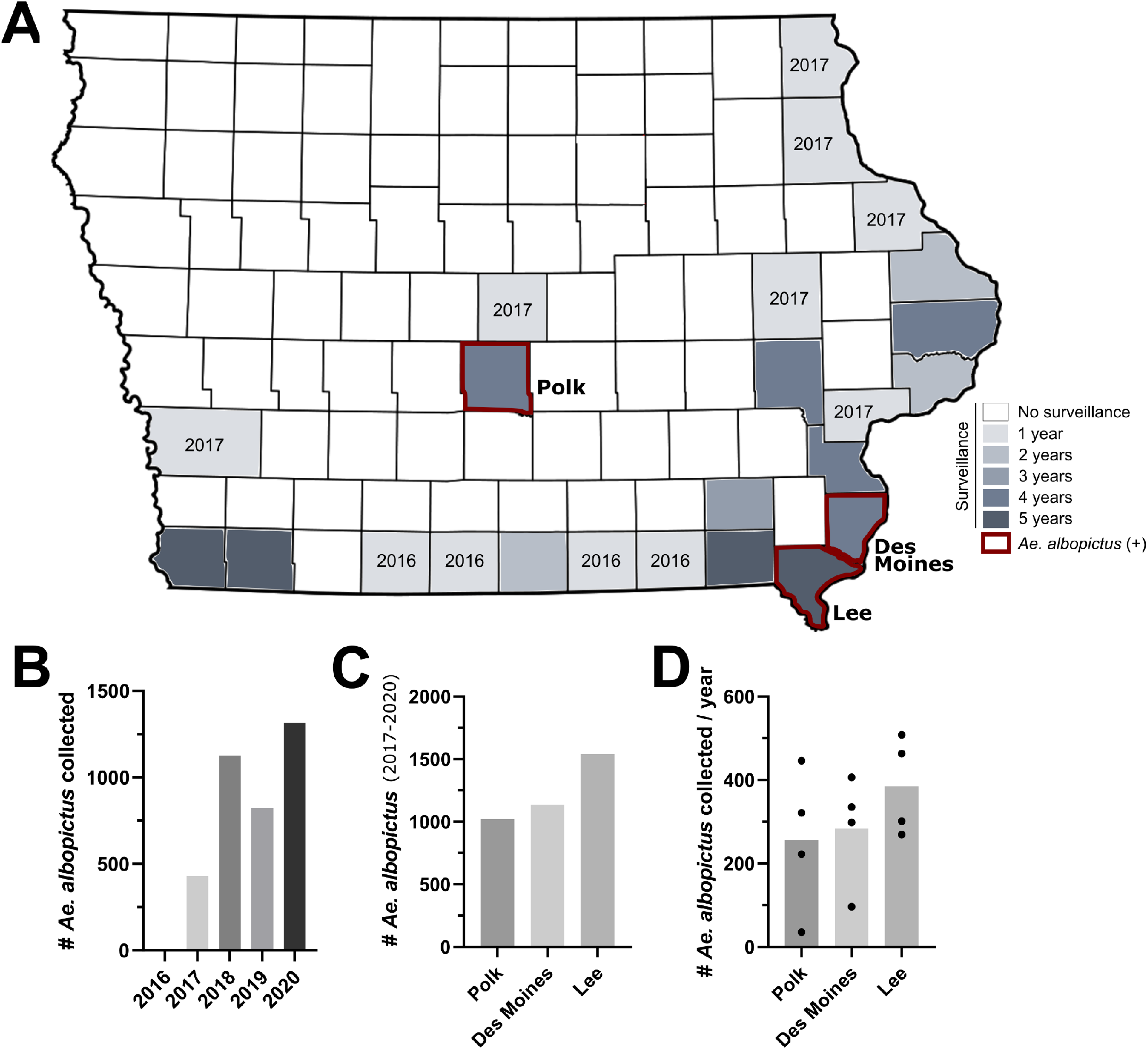

### *Ae. albopictus* population dynamics support their ability to overwinter in Iowa

To provide further support that *Ae. albopictus* have become established in each of the three counties, we examined *Ae. albopictus* weekly numbers and overall contributions to trapping yields within each of the mosquito trapping seasons from 2016-2020 in counties for which *Ae. albopictus* were detected (Figure 2). Across counties, *Ae. albopictus* populations reached peak abundance in late summer (late August, early September; approximately weeks 35 and 36), followed by sharp declines in abundance by early October (week 40) (Figure 2). For Polk County in central Iowa, *Ae. albopictus* was first detected in week 31 (early August) in 2017, yet in subsequent years (2018-2020) were consistently identified in mid-June (weeks 24 and 25; Figure 2A). Moreover, *Ae. albopictus* represented a much larger percentage of overall trap yields between 2018-2020 when compared to 2017 (Figure 2B), suggesting that their earlier abundance and higher proportion in the collected samples are indicative of their potential establishment.

**Figure 2.**
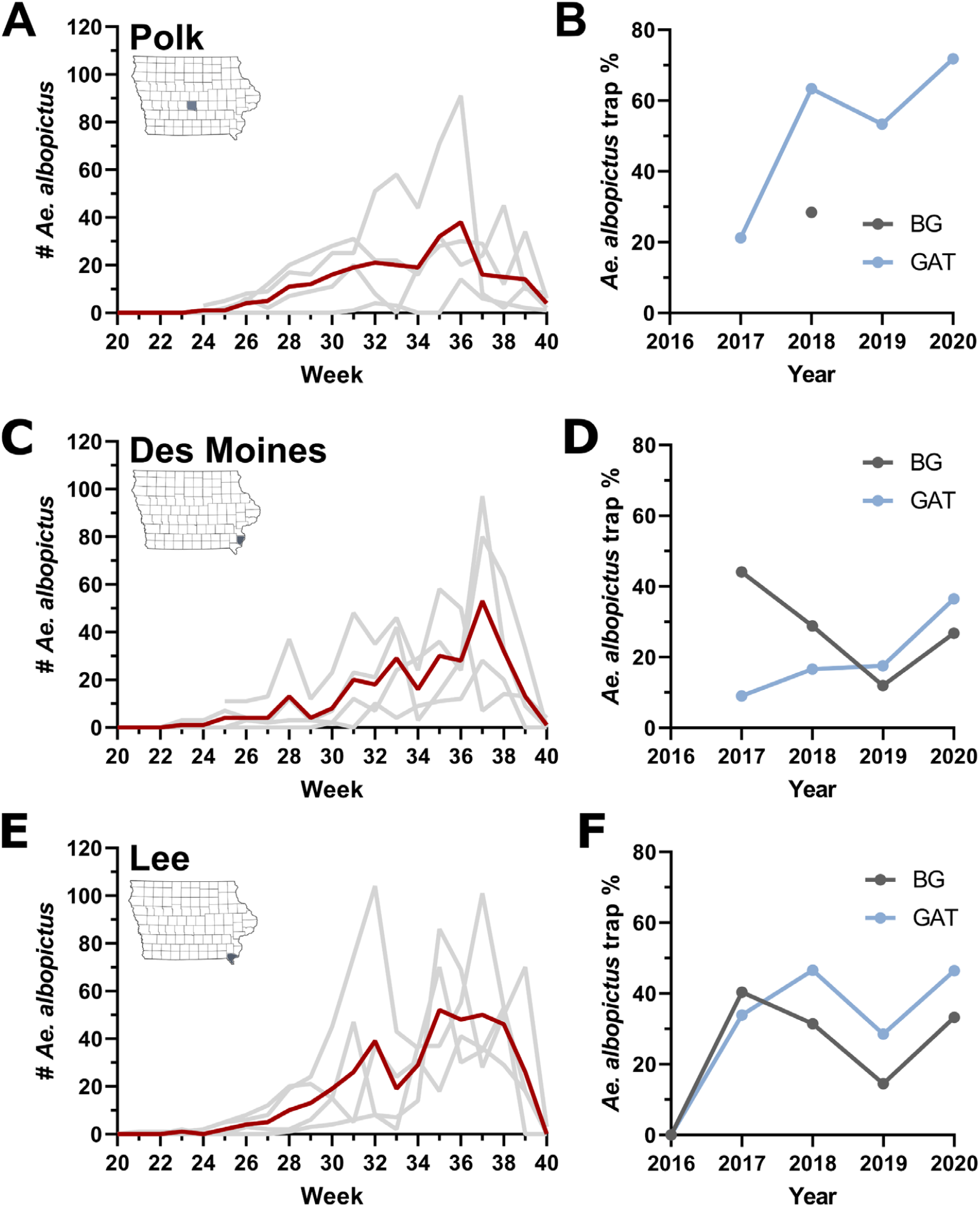
Abundance of *Ae. albopictus* in each positive Iowa county. The temporal abundance and percentage of *Ae. albopictus* in overall trap yields is displayed for Polk (**A, B**), Des Moines (**C, D**) and Lee County (**E, F**). Temporal abundance for each county (**A, C, E**) is displayed by epidemiological week, with the average abundance (2017-2020) displayed in red, while individual years are denoted by light gray lines. The percentage of *Ae. albopictus* collected of the total trap yields (**B, D, F**) are displayed for BG Sentinel (BG) and Gravid *Aedes* traps (GAT) for each year.

Similar patterns of *Ae. albopictus* abundance were also recorded in the southeastern portion of the state in Des Moines and Lee counties (Figure 2). For Des Moines County, the first detection of *Ae. albopictus* in 2017 occurred in week 30 (late July), while they were regularly detected in June (weeks 23-26) in subsequent years (2017-2020; Figure 2C). Although sites in Des Moines County displayed some variability between trap types, *Ae. albopictus* comprised between ∼10-40% of the overall number of mosquitoes collected in the county (Figure 2D). Lee County recorded the earliest detection of *Ae. albopictus* in 2017, with the first samples identified in week 28 (mid-July). In the years following (2018-2020), *Ae. albopictus* were found as early as week 21 (late-May; Figure 2E). Aside from 2016 when *Ae. albopictus* were not detected in our initial trapping efforts [19], *Ae. albopictus* represented ∼34% of the total trap yields when averaged across years and trap types (Figure 2F). While we account for some yearly variation in occurrence and overall abundance, these data provide further support for the overwintering and establishment of *Ae. albopictus* in multiple Iowa counties.

### Influence of landscape ecology on *Ae. albopictus* abundance

To determine if landscape ecology influences the presence of *Ae. albopictus* in Iowa, we looked to more closely examine the trapping site locations in each of the *Ae. albopictus-* positive counties. Trapping efforts in Polk County consisted of only a single site in close proximity to a facility involved in tire transport, with surrounding areas serving as an ideal habitat for *Ae. albopictus* (abundant breeding sites, tree cover, access to diverse hosts; Figure S1). Additional focused trapping efforts (BGs, GATs) were not performed at other locations in Polk County during this study. However, non-targeted surveillance involved with our West Nile virus surveillance program using other trap types (New Jersey Light Traps and Frommer Updraft Gravid Traps) have detected low numbers of *Ae. albopictus* in 2019 and 2020 at nearby locations in Polk County (Figure S2). The detection of *Ae. albopictus* in these suburban environments at locations that have been continuously trapped since 2016, suggests that *Ae. albopictus* have likely dispersed greater than 3 miles from their presumed point of introduction in recent years, providing further support for the ability of *Ae. albopictus* to overwinter in Polk County.

In contrast to the likely introduction of *Ae. albopictus* in Polk County associated with tire transport, there were no obvious mechanisms for the introduction of *Ae. albopictus* in Des Moines and Lee counties. As a result, the multiple trapping locations in Des Moines and Lee counties provided a better opportunity to determine the influence of landscape ecology on the presence or absence of *Ae. albopictus* (Figure 3). To address this question, we performed comparative land cover analysis on a total of 37 trapping site locations for which we defined each site for the presence/absence of *Ae. Albopictus* (Figure 3, Table S3). A total of 22 sites where *Ae. albopictus* were detected every year were considered “positive”, while the six sites for which *Ae. albopictus* were never found were considered “negative” (Table S3). Nine other sites where *Ae. albopictus* were identified but not collected every year were defined as “detected” (Table S3), suggesting that these sites represent new introductions that may or may not support established populations. Each of these trapping sites were mapped to their respective locations in Des Moines and Lee counties (Figure 3A), and the landscape ecology of “positive”, “negative”, and “detected” sites were compared (Figure 3B). We identified that areas of low-density development were significantly correlated with the presence of *Ae. albopictus*, while the percentage of agricultural areas were negatively associated with the presence of *Ae. albopictus* (Figure 3B). These data are supported by the spatial locations of the trapping site locations, where sites within urbanized areas were predominantly positive, while those located in more rural areas were typically negative (Figure 3A). This corresponds with the preferred habitat of *Ae. albopictus* which is most commonly associated with urban and suburban environments [37,38].

**Figure 3.**
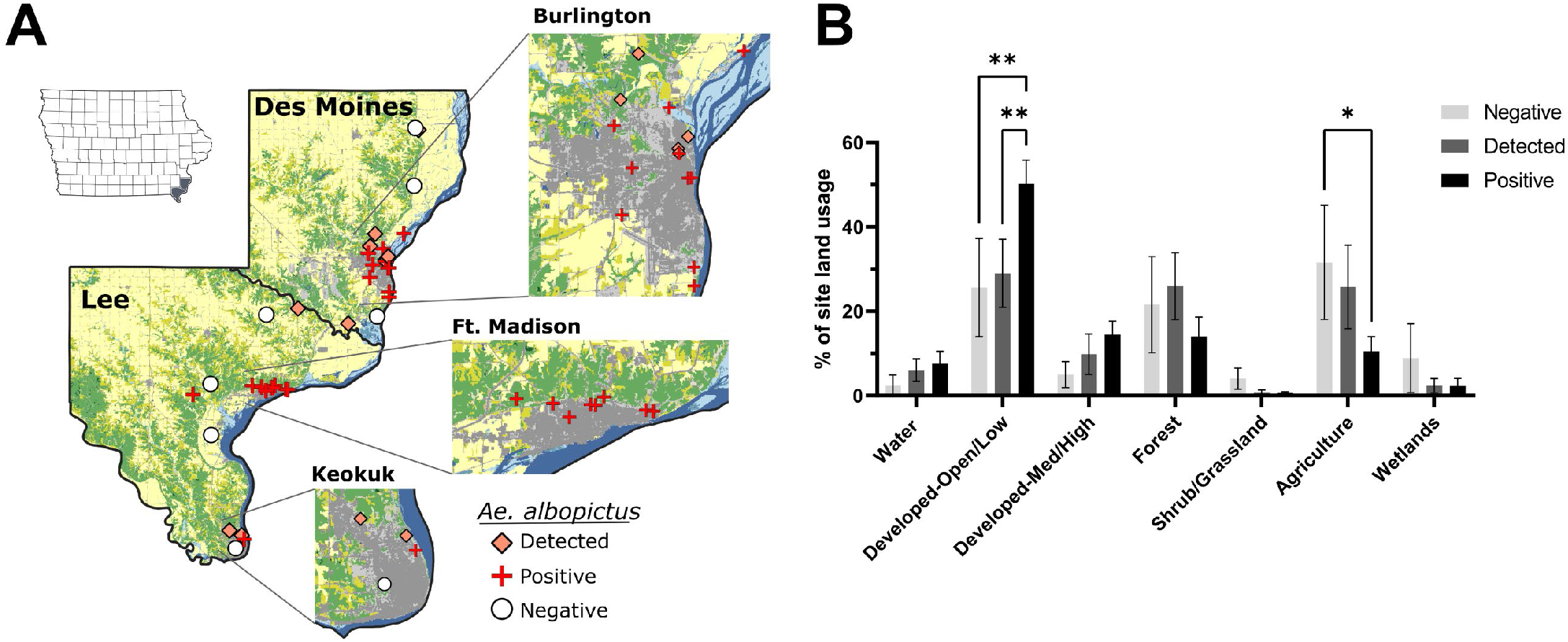
Landscape ecology influences *Ae. albopictus* abundance. Trapping site locations for both Des Moines and Lee counties display the presence/absence of *Ae. albopictus* as either positive (detected every year; red cross), detected (detected in some years; orange diamond), or negative (never detected; white circle) (**A**). Insets display expanded views for the most populous cities in Des Moines (Burlington) and Lee County (Ft. Madison and Keokuk). To better understand differences in the ecology of sites where *Ae. albopictus* were positive, detected, or negative, land use/land cover analysis was performed using 500m radius around each trapping location and displayed for different land use/land cover classifications (**B**). Statistical analysis was performed using a 2-way ANOVA and a Tukey’s multiple comparison test using GraphPad Prism software. Asterisks denote significance (*p < 0.05; **p < 0.01).

Furthermore, our trapping data provide support for the expansion of *Ae. albopictus* in both Des Moines and Lee counties. The presence of several sites for which *Ae. albopictus*, were detected, but not necessarily established, supports the movement of this mosquito species into new areas. This is further evidenced by the presence of *Ae. albopictus* in Keokuk (Lee County; Figure 3A) in 2018, where previous trapping efforts in 2016 [19] and 2017 suggested that these mosquito species were noticeably absent. This contrasts surveillance in Ft. Madison (Lee County; Figure 3A) where *Ae. albopictus* have been detected at every site since 2017 (Figure 3A, Table S3). With these two cities separated by ∼16 miles, these data imply that the distribution of *Ae. albopictus* in Lee County may be continually expanding into new locations.

### Genetic haplotype analysis identifies likely sources of introduction and supports overwintering of *Ae. albopictus* in Iowa

To better understand the origins of the *Ae. albopictus* collected in each of the three positive counties, DNA was isolated from a total of 165 individual samples and sequences of the mitochondrial CO1 gene were analyzed similarly to previous studies [16,33]. Sequence analysis resulted in the identification of 8 genetic haplotypes (Figure 4A, Table S4), distinguished by single nucleotide polymorphisms ranging between one to three nucleotides (Figure 4A, Table S5). The most abundant haplotypes, *hap_1* and *hap_3*, are distinguished by two nucleotides and were found in each of the three *Ae. albopictus* positive counties (Figure 4A). Both haplotypes represent common DNA haplotypes detected in other locations in the United States [16,33], southeast Asia [33,39], and Europe [33] (Table S4). Moreover, *hap_2* and *hap_3* were previously detected in Illinois [16] (Table S4), which due to its close proximity suggests that Illinois may serve as the likely origin and source for the introduction of *Ae. albopictus* in Iowa.

**Figure 4.**
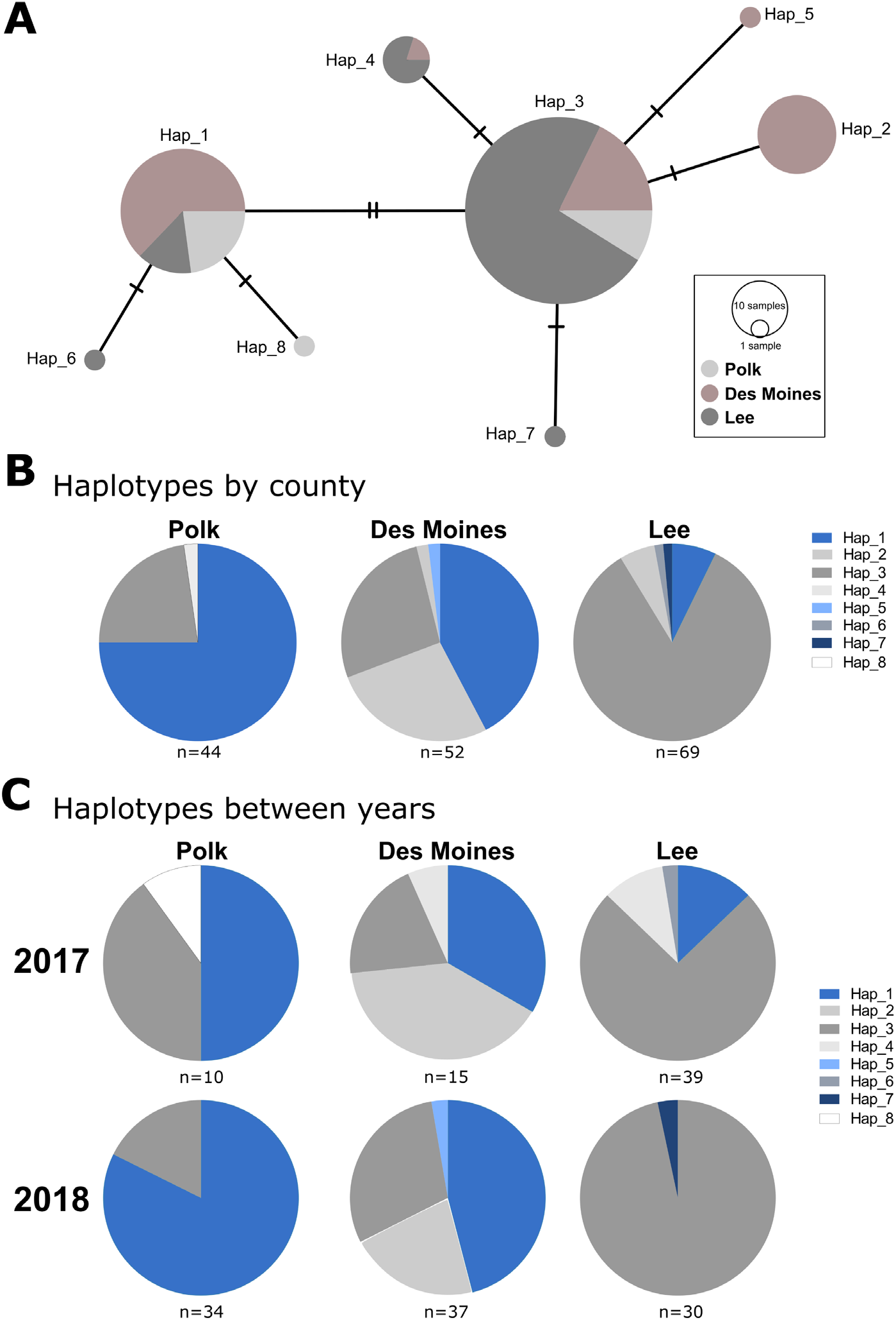
Comparative analysis of *Ae. albopictus* mtDNA haplotypes identified in Iowa. Comparisons of *Ae. albopictus* mtDNA haplotypes identified in Iowa are displayed as circles, with dashes on connecting lines indicating the number of nucleotide differences between haplotypes (**A**). Circle size corresponds to the number of individual samples, with differences in color representing the proportion samples from each respective county. Additional pie charts display differences in haplotypes between each Ae. albopictus- positive county (**B**) and the persistence of haplotypes between years (**C**).

For each of the Ae. albopictus counties, three or more haplotypes were detected, with both Des Moines and Lee counties displaying a total of five haplotypes (Figure 4B). While the majority of samples in Polk and Lee counties represented a single haplotype, Des Moines County displayed the most diverse population with *hap_1, 2*, and *3* comprising the majority of samples (Figure 4B). When samples were examined between years (2017 and 2018), the predominant haplotype(s) were consistent between years in each *Ae. albopictus* positive county (Figure 4C), providing further support for the establishment of these populations in each of the respective counties.

### *Ae. albopictus* overwintering in below freezing winter isotherms

Based on our surveillance data (Figure 2) and genetic analysis of DNA haplotypes (Figure 4), our results provide a strong argument for the introduction and establishment of *Ae. albopictus* in Iowa. Since overwintering temperatures have largely been attributed to limiting the spread of *Ae. albopictus* in North America [40], we examined the average winter temperature (December, January, February) isotherms for Iowa (1981-2010). Each of the *Ae. albopictus*-positive counties have average winter temperatures below freezing, with Des Moines and Lee counties in the -3 to -4°C isotherm, and Polk County in the -4 to -5°C isotherm (Figure 5A). While January temperatures (typically the coldest month) vary between years, temperatures ranged between -1 to -6°C in the *Ae. albopictus*-positive counties during our study period (Figure 5B). These low temperatures are traditionally considered to be not conducive to *Ae. albopictus* overwintering [18,40], suggesting that the *Ae. albopictus* populations in Iowa have been able to adapt to these freezing temperatures or have found adequate insulated environments to survive the winter. With the potential that global warming may have promoted elevated winter temperatures that may increase the chances of *Ae. albopictus* overwintering in the state, we examined the 30-year average January temperatures from 1981-2010 and 1991-2020 (Figure 5C). While Polk County displayed slightly warmer temperatures in recent years, temperatures in both Des Moines and Lee counties were ∼0.5°C cooler (Figure 5C), arguing that the recent expansion of Ae. albopictus into these areas is not the result of warmer winter temperatures.

**Figure 5.**
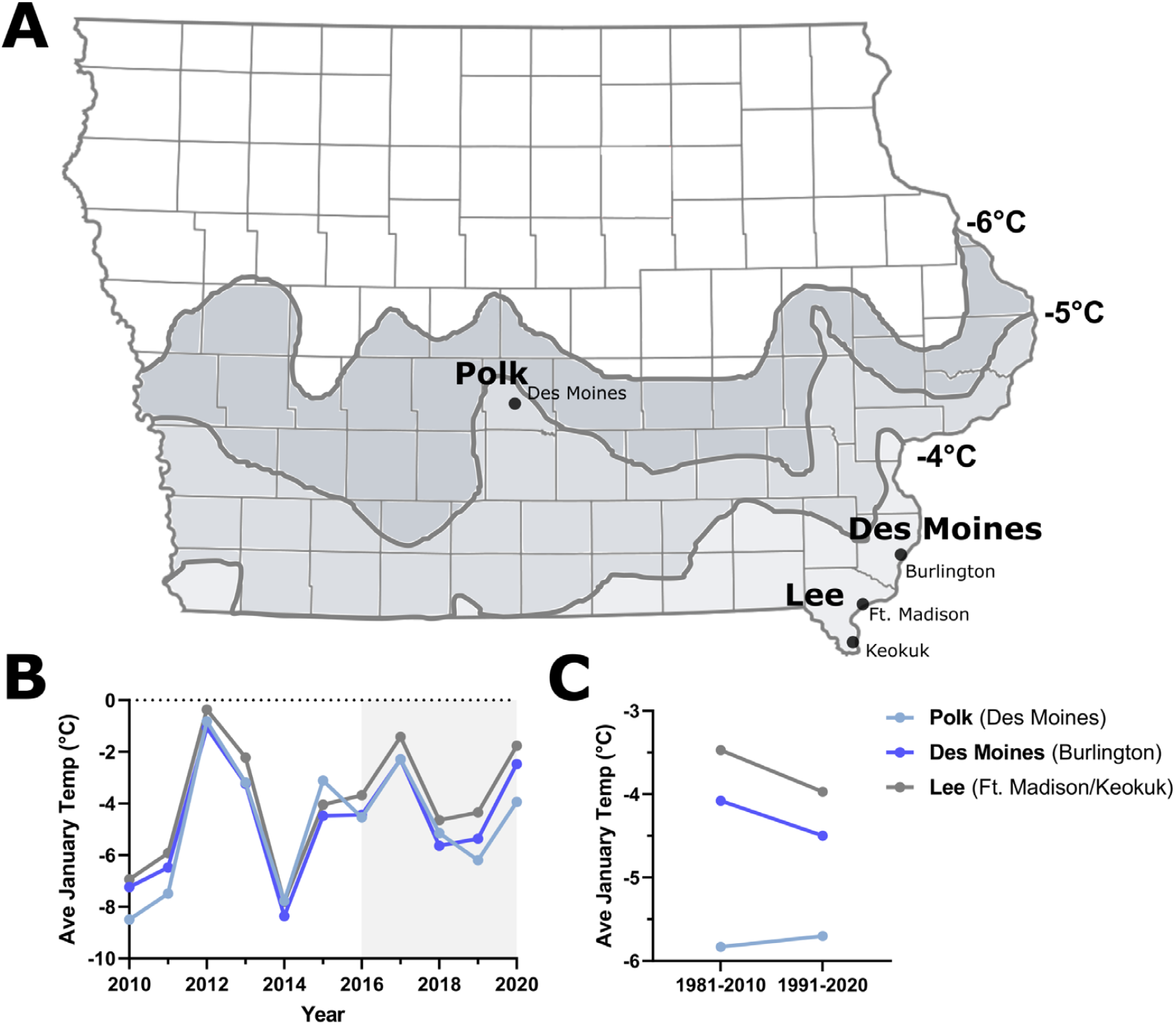
Overwintering temperatures in Iowa. Average winter (December, January, February) temperatures are displayed for Iowa (**A**). Shaded regions represent different temperature isotherms. Cities and counties where stable populations of *Ae. albopictus* have been detected are shown. Annual January temperatures are displayed from 2010-2020 to indicate differences in yearly temperatures for each respective *Ae. albopictus*-positive county (**B**). The shaded region from 2016-2020 represent the study period where targeted trapping efforts have focused on invasive *Aedes* species. Differences in January temperatures from the 30-year average from 1981-2010 and 1991-2020 examine the potential impacts of climate change on overwintering temperatures (**C**).

## Discussion

While limited detections of *Ae. albopictus* in Iowa (12 total from 1999-2016) have previously been described [17,41], these rare incidents were likely the result of isolated introduction events. However, with the introduction of Zika virus in the Americas in 2015-2016, there was an increased need to define the range of competent *Aedes* vectors throughout the US. Although our initial efforts in 2016 along the southern Iowa border did not detect *Ae. albopictus* [19], the results of our expanded surveillance efforts from 2017-2020 presented here describe the detection of more than 3,700 *Ae. albopictus* samples from three Iowa counties.

From these data, several lines of evidence support the establishment of *Ae. albopictus* in Iowa. This includes the consistent, early-season detection of *Ae. albopictus* in May and June, as well as the high percentage of the overall trapping yields for each of the three *Ae. albopictus*-positive counties. This is further validated by the occurrence of consistent mtDNA haplotypes between years, indicative of stable, genetic populations of *Ae. albopictus* that support mosquito overwintering.

While we cannot fully eliminate the possibility that new, yearly introductions may also contribute to the *Ae. albopictus* samples that were collected, the high number of individual mosquito samples collected for each *Ae. albopictus*-positive county makes this possibility unlikely. Although the primary site examined in Polk County is associated with tire transport, the surrounding areas are ideal *Ae. albopictus* habitat, with adequate tree cover, the presence of human and mammalian hosts, and an abundance of human-derived containers/tires that can serve as sites for oviposition that would support *Ae. albopictus* in much greater density. Although no obvious mechanisms of introduction by tire transport have been determined for Des Moines and Lee counties, the proximity to the Mississippi River and interstate highways that support human-associated transport likely account for the large number of *Ae. albopictus* collected in both locations. However, the consistency between years and the representation of similar *Ae. albopictus* mtDNA haplotypes across each county suggest that these are stable populations, supporting that the overwintering and establishment of *Ae. albopictus* as the most likely cause for the mosquitoes collected during our study period.

The presumed establishment of *Ae. albopictus* in Iowa challenges previous studies that have suggested that *Ae. albopictus* populations rarely extend north of 40°N latitude [7,18], presumably due to freezing winter temperatures that limit the ability of *Ae. albopictus* to overwinter in North America [40,42]. With average winter temperatures below 0°C which have traditionally limited overwintering and the expansion of *Ae. albopictus* [42], the freezing winter conditions in Iowa have traditionally been viewed as a major limitation to sustaining overwintering populations [18,40]. However, the detection of stable populations of *Ae. albopictus* in regions of Iowa with average winter temperature ranging from -3 to -5°C argue that these mosquito populations have potentially adapted to survive these harsh winter conditions. This also raises questions if winter temperatures alone are responsible for this climactic barrier, where the presence of snow cover may also help insulate overwintering *Ae. albopictus* eggs to enhance their survival [43]. Therefore, the microclimate of overwintering eggs may have a larger influence on the overwintering survival of *Ae. albopictus*.

An additional consideration of winter temperatures is the importance of urban heat islands [44–46], which may provide differences in microclimate across an urban and suburban landscape that results in warmer winter temperatures [45,46], potentially improving *Ae. albopictus* overwintering survival [47]. This is supported by the importance of urbanized development on the consistent detection of *Ae. albopictus* in Des Moines and Lee counties, where the presence of more rural, agricultural environments had significant influence on the presence/absence of *Ae. albopictus*. Furthermore, our site in Polk County also resides in an urbanized environment. Therefore, these urban microenvironments may provide increased “insulation” from the harsh winter temperatures that have been traditionally believed to serve as an ecological barrier for *Ae. albopictus* expansion. As a result, this would explain the higher density of *Ae. albopictus* collected in urban environments, and the low density or absence of *Ae. albopictus* collected in more natural or agricultural areas.

At present, it is unclear as to the exact timing of when *Ae. albopictus* were introduced into the state. Prior to 2016, mosquito surveillance in Iowa predominantly focused on West Nile virus [31], utilizing a trap network that was not ideal for the collection of invasive *Aedes* species and did not extend into areas that would most likely serve as points of introduction. In 2016, our initial surveillance efforts along Iowa’s southern border failed to detect *Ae. albopictus* [19], yet in hindsight, these predominantly agricultural and less densely populated areas did not represent ideal *Ae. albopictus* habitats. However, our trapping efforts in 2016 did include Lee County [19], where our more focused efforts described in this study from 2017-2020 did result in the detection of established populations of *Ae. albopictus*. This discrepancy is most likely due to differences in trap locations in Lee County when compared between 2016 and 2017-2020, which for 2016 included only sites near Keokuk, while in 2017-2020 included both Keokuk and Ft. Madison. It is therefore of note that beginning in 2017 and in subsequent years, each of the trapping sites in Ft. Madison were positive for *Ae. albopictus*. Yet, for Keokuk which is ∼16 miles away, the first detection of *Ae. albopictus* wasn’t until 2018. This includes at least one site with continual trapping efforts in 2016 [19] and 2017 for which *Ae. albopictus* was later detected in 2018. These intra-county differences suggest that the introduction of *Ae. albopictus* in Lee County likely occurred prior to 2017 for Ft. Madison, while the introduction into Keokuk is more recent, potentially even during the years of our study (2017-2020) and suggests that their distribution is continuing to expand in the county. Similar to Ft. Madison (Lee County), we believe that the detection of *Ae. albopictus* in Burlington (Des Moines County) occurred prior to our trapping efforts in 2017 based on their abundance and distribution throughout the city. However, previous surveillance efforts in Des Moines County have been limited, preventing the determination of a definitive timeline for their introduction. Based on the prevalence of multiple mtDNA haplotypes in Des Moines County, multiple invasion events may have contributed to the established *Ae. albopictus* populations in the county. For both Des Moines and Lee counties, the proximity of the Mississippi River and interstate highways to the urban areas of both counties likely represents the most feasible route of introduction through freight and shipping along the waterway or from interstate transport from neighboring Illinois.

In contrast, the introduction of *Ae. albopictus* into Polk County, are inextricably tied to the tire transport industry. Due to the potential for yearly infestations, which may be responsible for previous detections of *Ae. albopictus* in the county [7,8,17], is difficult to definitively demonstrate that the populations of *Ae. albopictus* identified in Polk County are of established populations and not an annual infestation. Yet, during the course of our study (2017-2020), there is strong evidence that we may have captured a local infestation that was able to overwinter and establish in the area. Support for this includes an ∼10-fold increase in the number of *Ae. albopictus* collected between 2017 and 2018 (35 and 321 respectively), a dramatic increase in the percentage of *Ae. albopictus* in overall gravid *Aedes* trap yields (21% in 2017 compared to an average of 62% from 2018-2020), and the consistent detection of two predominant mtDNA haplotypes (*hap_1* and *hap_3*) between 2017 and 2018. Moreover, the recent detection of *Ae. albopictus* at low densities at other non-focused trapping sites in close proximity to the tire facility support their potential expansion and establishment in the area.

Together, our results provide strong evidence for the presence and establishment of *Ae. albopictus* populations in Iowa, demonstrating the further expansion of *Ae. albopictus* into the Upper Midwest region of the United States. Importantly, with consistent winter temperatures in Iowa that are below freezing, this challenges existing beliefs that these winter temperature extremes serve as the primary boundary for *Ae. albopictus* overwintering and expansion [18,39,40,42]. With the additional recent detection of *Ae. albopictus* in Wisconsin [15], this raises an increased need for continual surveillance to monitor the further spread and expansion of *Ae. albopictus* in the Upper Midwest and other regions of the world on the northern range of the expansion of *Ae. albopictus*. Through increased urbanization and predicted climate change, the distribution of *Ae. albopictus* is only expected to further spread [3], highlighting the increased risk of mosquito-borne disease transmission in new regions of the world.

## Acknowledgements

We would like to thank Julie Coughlin of the Iowa Department of Public Health, the many local public health partners that contributed to our mosquito trapping efforts, especially to Christa Poggemiller of the Des Moines County Public Health Department and Michele Ross of the Lee County Health Department for leading these efforts. We would also like to thank those that enabled access to their properties to conduct mosquito trapping, Chris Lee for assistance in mosquito identifications, and to Chris Stone for discussions regarding the DNA haplotype analysis. This research was supported by the USDA National Institute of Food and Agriculture, Hatch Project 101071, the Epidemiology and Laboratory Capacity for Infectious Diseases (ELC) Program, and the Midwest Center of Excellence for Vector-Borne Disease. This publication was supported by Cooperative Agreement #U01 CK000505, funded by the Centers for Disease Control and Prevention. Its contents are solely the responsibility of the authors and do not necessarily represent the official views of the Centers of Disease Control and Prevention or the Department of Health and Human Services.

## References

1. Reiter P, Sprenger D. The used tire trade: a mechanism for the worldwide dispersal of container breeding mosquitoes. J Am Mosq Control Assoc. 1987;3: 494–501.

2. Kraemer MUG, Sinka ME, Duda KA, Mylne A, Shearer FM, Brady OJ, et al. The global compendium of Aedes aegypti and Ae. albopictus occurrence. Sci Data. 2015;2: 150035.

3. Kraemer M, Reiner R, Brady O, Messina J, Gilbert M, Pigott D, et al. Past and future spread of the arbovirus vectors Aedes aegypti and Aedes albopictus. Nat Microbiol. 2019;4: 854–863.

4. Bonizzoni M, Gasperi G, Chen X, James AA. The invasive mosquito species Aedes albopictus: Current knowledge and future perspectives. Trends Parasitol. 2013;29: 460–468.

5. Sprenger D, Wuithiranyagool T. The discovery and distribution of Aedes albopictus in Harris County, Texas. J Am Mosq Control Assoc. 1986;2: 217–219.

6. Yee DA. Thirty years of Aedes albopictus (Diptera: Culicidae) in America: An introduction to current perspectives and future challenges. J Med Entomol. 2016;53: 989–991.

7. Hahn MB, Eisen RJ, Eisen L, Boegler KA, Moore CG, McAllister J, et al. Reported distribution of Aedes (Stegomyia) aegypti and Aedes (Stegomyia) albopictus in the United States, 1995-2016 (Diptera: Culicidae). J Med Entomol. 2016;53: 1169– 1175.

8. Hahn MB, Eisen L, McAllister J, Savage HM, Mutebi J, Eisen RJ. Updated reported distribution of Aedes (Stegomyia) aegypti and Aedes (Stegomyia) albopictus (Diptera : Culicidae) in the United States, 1995 – 2016. 2017;54: 1420–1424.

9. Egizi A, Healy SP, Fonseca DM. Rapid blood meal scoring in anthropophilic Aedes albopictus and application of PCR blocking to avoid pseudogenes. Infect Genet Evol. 2013;16: 122–128.

10. Paupy C, Delatte H, Bagny L, Corbel V, Fontenille D. Aedes albopictus, an arbovirus vector: From the darkness to the light. Microbes Infect. 2009;11: 1177– 1185.

11. Grard G, Caron M, Mombo IM, Nkoghe D, Mboui Ondo S, Jiolle D, et al. Zika Virus in Gabon (Central Africa) - 2007: A New Threat from Aedes albopictus? PLoS Negl Trop Dis. 2014;8: e2681.

12. McKenzie BA, Wilson AE, Zohdy S. Aedes albopictus is a competent vector of Zika virus: A meta-analysis. PLoS One. 2019;14: e0216794.

13. Claborn DM, Poiry M, Famutimi OD, Duitsman D, Thompson KR. A survey of mosquitoes in southern and western missouri. J Am Mosq Control Assoc. 2018;34: 131–133.

14. Janousek TE, Plagge J, Kramer WL. Record of Aedes albopictus in Nebraska with notes on its biology. J Am Mosq Cont Control Assoc. 2001;17: 265–267.

15. Richards T, Tucker BJ, Hassan H, Bron GM, Bartholomay L, Paskewitz S. First detection of Aedes albopictus (Diptera: Culicidae) and expansion of Aedes japonicus japonicus in Wisconsin, United States. J Med Entomol. 2019; 56:291–296.

16. Stone CM, Zuo Z, Li B, Ruiz M, Swanson J, Hunt J, et al. Spatial, temporal, and genetic invasion dynamics of Aedes albopictus (Diptera: Culicidae) in Illinois. J Med Entomol. 2020;57: 1488–1500.

17. Dunphy BM, Rowley WA, Bartholomay LC. A taxonomic checklist of the mosquitoes of Iowa. J Am Mosq Control Assoc. 2014;30: 119–121.

18. Johnson TL, Haque U, Monaghan AJ, Eisen L, Hahn MB, Hayden MH, et al. Modeling the environmental suitability for Aedes (Stegomyia) aegypti and Aedes (Stegomyia) albopictus (Diptera: Culicidae) in the contiguous United States. 2017; J Med Entomol. 2017;54: 1605–1614.

19. Kovach KB, Smith RC. Surveillance of mosquitoes (Diptera: Culicidae) in southern Iowa, 2016. J Med Entomol. 2018;55: 1341–1345.

20. Eiras AE, Buhagiar TS, Ritchie SA. Development of the Gravid Aedes Trap for the capture of adult female container-exploiting mosquitoes (Diptera: Culicidae). J Med Entomol. 2014;51: 200–209.

21. Maciel-de-Freitas R, Eiras ÁE, Lourenço-de-Oliveira R. Field evaluation of effectiveness of the BG-Sentinel, a new trap for capturing adult Aedes aegypti (Diptera: Culicidae). Mem Inst Oswaldo Cruz. 2006;101: 321–325.

22. Farajollahi A, Kesavaraju B, Price DC, Williams GM, Healy SP, Gaugler R, et al. Field efficacy of BG-Sentinel and industry-standard traps for Aedes albopictus (Diptera: Culicidae) and West Nile virus surveillance. J Med Entomol. 2009;46: 919–25.

23. Meeraus WH, Armistead JS, Arias JR. Field comparison of novel and gold standard traps for collecting Aedes albopictus in Northern Virginia. J Am Mosq Control Assoc. 2008;24: 244–248.

24. Johnson BJ, Hurst T, Quoc HL, Unlu I, Freebairn C, Faraji A, et al. Field comparisons of the Gravid Aedes Trap (GAT) and BG-Sentinel Trap for monitoring Aedes albopictus (Diptera: Culicidae) populations and notes on indoor GAT collections in Vietnam. J Med Entomol. 2018;54: 340–348.

25. Darsie R, Ward R. Identification and geographical distribution of the mosquitoes of North America, North of Mexico. University Press of Florida; 2005.

26. Multi-Resolution Land Characteristics Consortium. NLCD 2016 Land Cover (CONUS).

27. Bonnet DD, Worcester DJ. The dispersal of Aedes albopictus in the territory of Hawaii. Am J Trop Med Hyg. 1946;26: 465–476.

28. Niebylski ML, Craig GB. Dispersal and survival of Aedes albopictus at a scrap tire yard in Missouri. J Am Mosq Control Assoc. 1994;10: 339–343.

29. Verdonschot PFM, Besse-Lototskaya AA. Flight distance of mosquitoes (Culicidae): A metadata analysis to support the management of barrier zones around rewetted and newly constructed wetlands. Limnologica. 2014;45: 69–79.

30. Post RJ, Flook PK, Millest AL. Methods for the preservation of insects for DNA studies. Biochem Syst Ecol. 1993;21: 85–92.

31. Dunphy BM, Kovach KB, Gehrke EJ, Field EN, Rowley WA, Bartholomay LC, et al. Long-term surveillance defines spatial and temporal patterns implicating Culex tarsalis as the primary vector of West Nile virus. Sci Rep. 2019;9: 6637.

32. Field EN, Gehrke EJ, Ruden RM, Adelman JS, Smith RC. An improved multiplex Polymerase Chain Reaction (PCR) assay for the identification of mosquito (Diptera: Culicidae) blood meals. J Med Entomol. 2020;57: 557–562.

33. Zhong D, Lo E, Hu R, Metzger ME, Cummings R, Bonizzoni M, et al. Genetic analysis of invasive Aedes albopictus populations in Los Angeles County, California and its potential public health impact. PLoS One. 2013;8: e68586.

34. Rozas J, Ferrer-Mata A, Sanchez-DelBarrio JC, Guirao-Rico S, Librado P, Ramos-Onsins SE, et al. DnaSP 6: DNA sequence polymorphism analysis of large data sets. Mol Biol Evol. 2017;34: 3299–3302.

35. Leigh J, Bryant D. POPART: full-feature software for haplotype network construction. Methods Ecol. Evol. 2015;6: 1110–1116.

36. Ogden R, Shuttleworth C, McEwing R, Cesarini S. Median-joining networks for inferring intraspecific phylogenies. Conserv Genet. 2005;6: 37–48.

37. Braks MAH, Honório NA, Lourenço-De-Oliveira R, Juliano SA, Lounibos LP. Convergent habitat segregation of Aedes aegypti and Aedes albopictus (Diptera: Culicidae) in southeastern Brazil and Florida. J Med Entomol. 2003;40: 785–794.

38. Delatte H, Toty C, Boyer S, Bouetard A, Bastien F, Fontenille D. Evidence of habitat structuring Aedes albopictus populations in Réunion Island. PLoS Negl Trop Dis. 2013;7: e2111.

39. Lee EJ, Yang SC, Kim TK, Noh BE, Lee HS, Kim H, et al. Geographical Genetic variation and sources of Korean Aedes albopictus (Diptera: Culicidae) populations. J Med Entomol. 2020;57: 1057–1068.

40. Nawrocki SJ, Hawley WA. Estimation of the northern limits of distribution of Aedes albopictus in North America. J Am Mosq Control Assoc. 1987;3: 314–317.

41. Moore CG. Aedes albopictus in the United States: Current status and prospects for further spread. J Am Mosq Control Assoc. 1999;15: 221–227.

42. Armstrong PM, Andreadis TG, Shepard JJ, Thomas MC. Northern range expansion of the Asian tiger mosquito (Aedes albopictus): Analysis of mosquito data from Connecticut, USA. PLoS Negl Trop Dis. 2017;11: e0005623.

43. Rochlin I, Ninivaggi D V., Hutchinson ML, Farajollahi A. Climate change and range expansion of the Asian tiger mosquito (Aedes albopictus) in Northeastern USA: Implications for public health practitioners. PLoS One. 2013;8: e60874.

44. Zhao L, Lee X, Smith RB, Oleson K. Strong contributions of local background climate to urban heat islands. Nature. 2014;511: 216–219.

45. Yang J, Bou-Zeid E. Should cities embrace their heat islands as shields from extreme cold? J Appl Meteorol Climatol. 2018;57: 1309–1320.

46. Macintyre HL, Heaviside C, Cai X, Phalkey R. Comparing temperature-related mortality impacts of cool roofs in winter and summer in a highly urbanized European region for present and future climate. Environ Int. 2021;154.

47. Ward TB. Influence of an urban heat island on mosquito development and survey of biting midge species associated with white-tailed deer farms. Oklahoma State University. 2011.

