## Supporting Figures S1 and S2 for "A tiger in the Upper Midwest: Surveillance and genetic data support the introduction and establishment of *Aedes albopictus* in Iowa, USA"

[2]Current address: Department of International and Global Studies, Mercer University, Macon, Georgia, USA

### Supporting Information

### Supporting Figures

**Figure S1.** Surrounding areas adjacent to the site of introduction in Polk County represent ideal habitats for *Ae. albopictus*.

**Figure S2.** Spread of *Ae. albopictus* in suburban areas of Polk County.

### Supporting Tables

**Table S1.** Overview of Iowa counties participating in *Aedes* trapping efforts.

**Table S2.** Trapping efforts across counties (2017-2020).

**Table S3.** Land use analysis of present, absent, or detected sites for *Ae. albopictus*.

**Table S4.** Description of *Ae. albopictus* mtDNA haplotypes.

**Table S5.** Nucleotide polymorphisms defining *Ae. albopictus* mtDNA haplotypes.

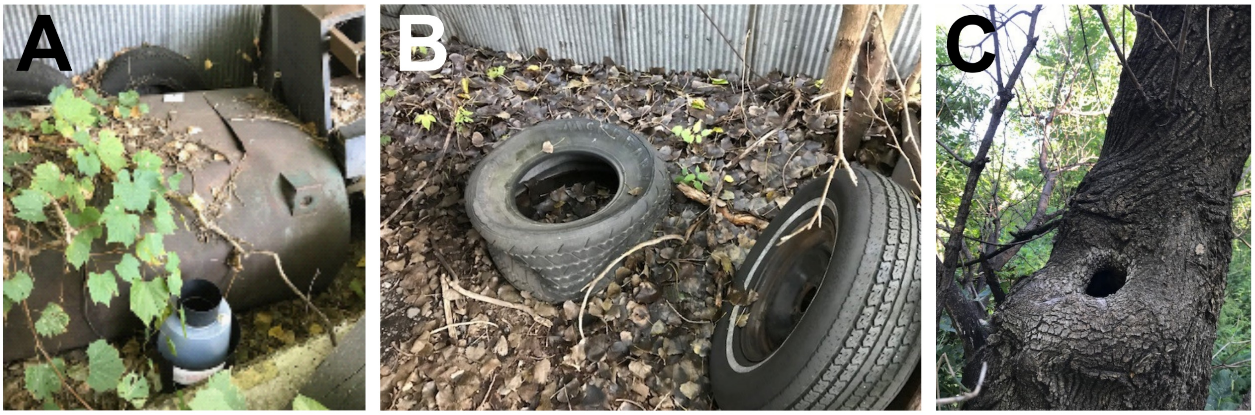


**Figure S1. Surrounding areas adjacent to the site of introduction in Polk County represent ideal habitats for *Ae. albopictus*.** The focal trapping location in Polk County is a neglected commercial property with ample vegetation, junk, and debris (**A**). This includes several used tires with adequate leaf litter (**B**) and tree holes (**C**) which can be used as potential oviposition sites for *Ae. albopictus*.

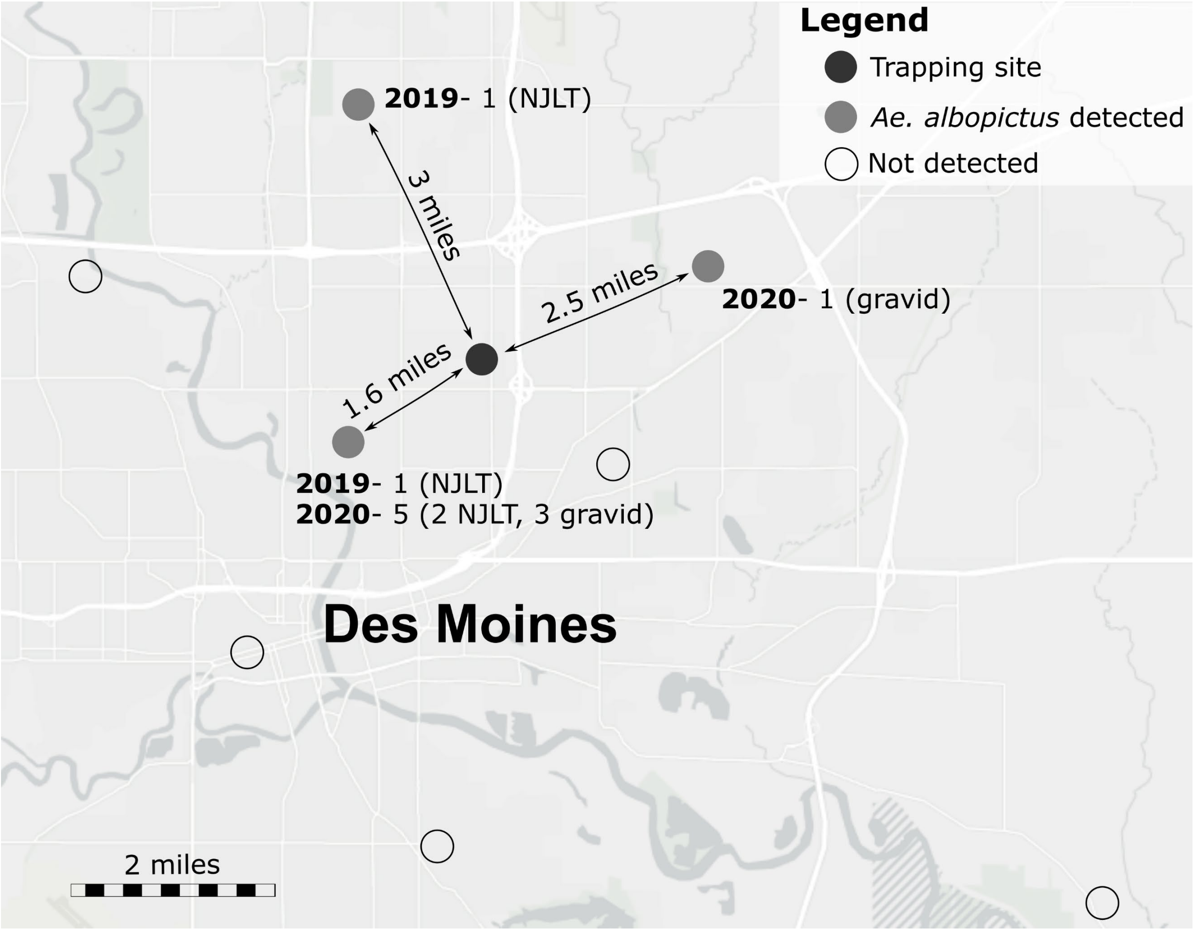


**Figure S2. Spread of *Ae. albopictus* in suburban areas of Polk County.** *Ae. albopictus* have been detected at additional trapping site locations in the greater Des Moines metropolitan area near the focal trapping location using either New Jersey Light Traps (NJLT) or Frommer Updraft Gravid Traps (gravid). Approximate distances between the primary site location (black dot) and other *Ae. albopictus*-positive sites (grey dots) are displayed. Additional trapping sites where *Ae. albopictus* have not been detected are shown with open circles.
